# *traceax*: a JAX-based framework for stochastic trace estimation

**DOI:** 10.1101/2025.07.14.662216

**Authors:** Abdullah Al Nahid, Linda Serafin, Nicholas Mancuso

## Abstract

In many applications, from statistical inference to machine learning, calculating the trace of a matrix is a fundamental operation, yet may be infeasible due to memory constraints. Stochastic trace estimation offers a practical solution by using randomized matrix-vector products to obtain accurate, unbiased estimates without constructing the full matrix in memory. Here, we present *traceax*, a Python framework for scalable trace estimation that leverages efficient linear operator representations of matrices while supporting automatic differentiation and hardware acceleration. *traceax* supports state-of-the-art trace estimators and through simulations we recapitulate results demonstrating their high accuracy while significantly reducing runtime and memory usage when compared with direct trace computation. As a proof of concept, we implemented a stochastic heritability estimator using traceax requiring only several lines of code. Overall, *traceax* provides a versatile tool for stochastic trace estimation that can be easily integrated into existing inferential pipelines.

*traceax* is freely available at: https://github.com/mancusolab/traceax

## INTRODUCTION

High dimensionality is the norm in modern statistical inference, machine learning, and computational biology, where data operators routinely involve matrices with *N* well into the millions. The trace of such operators remains essential for applications such as variance-component estimation, log-marginal likelihoods, and matrix-function quadrature (Cortinovis and Kressner, 2022; Ubaru et al., 2017; Ubaru and Saad, 2018; Yeon et al., 2025). However, explicitly forming or storing the full matrix can be impractical. Stochastic trace estimation offers an elegant workaround by sampling a small number of random probe vectors, performing matrix-vector products, and averaging the results. This approach produces an unbiased estimate of tr(***A***) without ever fully materializing ***A***. Researchers over the past few decades have developed estimators such as Hutchinson, Hutch++, XTrace, XNysTrace (Epperly et al., 2024; Hutchinson, 1990; Meyer et al., 2021; Yeon et al., 2025; Zvonek et al., 2024); however, their implementations are often incompatible with automatic differentiation (autodiff) and do not take advantage of modern computing architectures.

To address these limitations, we propose *traceax*, a Python-based software that implements stochastic trace estimators with an easy-to-use API. It leverages the flexibility of linear operators (as implemented in *lineax* (Rader et al., 2023)) together with differentiable and performant JAX-based numerics. JAX has gained traction in performance-critical scientific domains by uniting NumPy-like syntax, autodiff, and just-in-time (JIT) compilation within a framework that scales efficiently across modern hardware (Bradbury et al., 2018; Gallup et al., 2024; Kaymak et al., 2022; Zhang et al., 2025).

Under simulations, we demonstrate our implementation of recent estimators in *traceax* recapitulates results from alternative non-JAX implementations (Epperly et al., 2024). Finally, as a proof of concept, we show that *traceax* can be straightforwardly applied to heritability estimation (Wu and Sankararaman, 2018), a fundamental operation in statistical genetics. Overall, our results demonstrate that *traceax* provides an intuitive and scalable framework for stochastic trace estimation that integrates seamlessly with modern scientific-computing workflows.

## BACKGROUND & IMPLEMENTATION

First, we briefly describe stochastic trace estimation. Namely, given matrix ***A*** ∈ ℝ^*N*×*N*^ the trace is defined as tr(***A***) = ∑_*i*_ ***A***_*ii*_. In practice, however, we often need to compute the trace of a matrix that is the result of some linear algebraic expression (e.g., ***A*** = ***XX***′). Thus, we consider ***A*** to be a linear operator, whose internal representation is abstracted, which precludes inspecting individual elements in ***A***, however, we are allowed to perform matrix-vector products ***Av*** for any ***v***. Hutchinson and others (Epperly et al., 2024; Hutchinson, 1990; Meyer et al., 2021) demonstrated that given *k* random probe vectors ***v***_(1)_, …, ***v***_(*k*)_, such that 𝔼[***vv***^⊤^] = ***I***_*N*_, an unbiased estimator is given by,

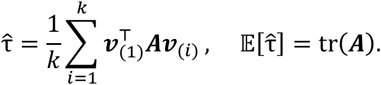

*traceax* automates this workflow and utilizes JIT compilation for highly performant numerics that can be run on CPU, GPU, or TPU by setting a single flag. It also supports autodiff due to JAX, which enables trace estimation as an intermediate operation in larger optimization procedures.

*traceax* currently supports four trace estimators, Hutchinson (Hutchinson, 1990), Hutch++ (Meyer et al., 2021), XTrace, and XNysTrace (Epperly et al., 2024). It allows sampling random probe vectors from Gaussian, Rademacher, and Spherical distributions, with reasonable defaults. For details on arguments and options, see the documentation.

To illustrate the simplicity and usability of our tool, we have provided example scripts in the repository that implement a scalable method-of-moment estimator to approximate SNP-heritability 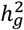 from large biobank-scale genetic data.

## APPLICATION

First, to demonstrate the correctness of our implemented estimators in *traceax*, we evaluated synthetic symmetric operators similar to the settings described in Epperly et al. (Epperly et al., 2024). Importantly, *traceax* recapitulated the relative performance described in Epperly et al. (Epperly et al., 2024). We found that all estimators are consistent as the mean relative error either decreases steadily or remains stable as *N* or *k* increases (**Figure 1**). Second, variance-reduction techniques paid off most when the spectrum was highly non-uniform. In both the polynomial and exponential settings, XTrace, and especially XNysTrace, reach sub-10^−5^ accuracy with as few as *k*=100 probes, whereas Hutchinson remained four to five orders of magnitude less accurate at the same budget.

**Figure 1:**
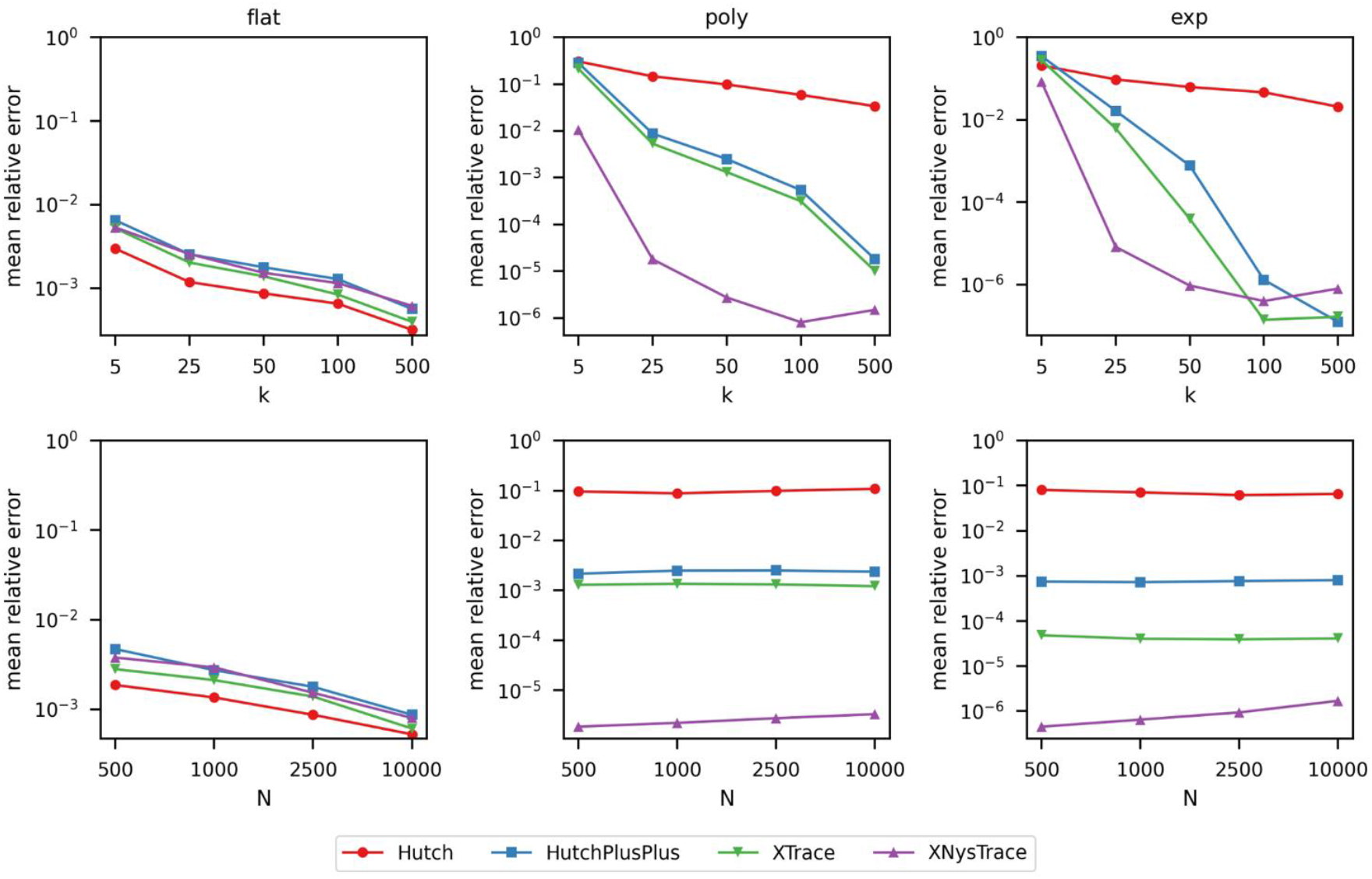
Mean relative error of stochastic trace estimators across spectral settings. *Top row:* mean relative error (log scale) plotted against probe size *k*, with fixed matrix size *N*=2500. *Bottom row:* mean relative error plotted against *N* with fixed *k*=50. Columns correspond to different eigenvalue spectra: flat (left), polynomial decay (middle), and exponential decay (right). The four methods compared are Hutch, Hutch++, XTrace, and XNysTrace. Each configuration was run 100 times with default parameters unless otherwise specified.

We next examined the effect of the probe distribution on trace estimation quality (**Figure S1**). We used the XTrace estimator in *traceax* with Gaussian, Rademacher, and Spherical probe distributions and measured the average relative error. We found that the choice of probe distribution does not confound accuracy as the results were consistent across all distributions.

To assess runtime performance, we measured the average execution times for all estimators in *traceax*. Runtime increased approximately linearly with the number of probes *k* (**Figure S2**). In the polynomial and exponential spectra, Hutchinson was the fastest, followed by Hutch++, then XTrace, with XNysTrace being the slowest. In contrast, in the flat-spectrum setting, Hutch++ was the fastest, while Hutchinson had the slowest runtime. Importantly, all estimators have an average runtime of under one second, which makes them well suited for rapid, exploratory workflows. In addition, we evaluated the performance gain from JIT compilation by running XTrace in both JIT and non-JIT settings (**Figure S3**). Unsurprisingly, JIT compilation accelerates every estimator by one to two orders of magnitude. However, this gain surprisingly decreases as *N* or *k* increases.

Finally, to demonstrate a realistic application, we reimplemented the Randomized Haseman-Elston regression (RHE-reg) framework for SNP-heritability estimation (Wu and Sankararaman, 2018). Briefly, heritability estimation aims to quantify the proportion of variance in an observed trait or phenotype that can be explained by additive genetic effects captured by genome-wide single nucleotide polymorphisms (SNPs). Using simulated genotype matrices (***X*** ∈ ℝ^*N*×*M*^) and phenotypes with true heritability 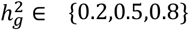, we estimated tr(***K***) and tr(***K***^2^) for 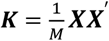 with varying *N* samples, *M* SNPs, and *k* random probes. We found that all estimators in *traceax* provide unbiased estimates of 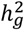 with inter-quartile ranges shrinking as the scale of data grows (**Figure S4**).

To compare the efficiency of *traceax* estimators, we then used the RHE-reg SNP-heritability estimator reimplementation in two settings: one using direct trace computation and the other using the XTrace-based stochastic trace estimator in *traceax*. We evaluated both approaches by measuring memory usage and runtime across 10 independent runs (**Figure S5**). On average, direct trace consumed nearly 2x the memory (∼38 GiB) and the runtime was about 30x longer (∼17 minutes) compared to the stochastic trace (∼19 GiB of memory and <1 minute). In both cases, 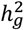 estimates were highly accurate (see **Supplementary Table**).

Collectively, these results demonstrate that *traceax* combines statistical efficiency with hardware acceleration, which makes large-scale stochastic trace estimation both accurate and practical.

## CONCLUSION

In this work, we developed *traceax*, a JAX-based Python library that implements four stochastic trace estimators through a simple and modular interface. Its design enables easy extension with custom estimators and probe distributions by extending abstract base classes.

Our results show that *traceax* produces accurate trace estimates with low runtime across diverse spectral conditions. These findings establish *traceax* as a scalable and extensible tool for stochastic trace estimation. By uniting performance, modularity, and ease of use, *traceax* offers a practical solution for future work in large-scale linear operator computation.

## Acknowledgments

We thank members of Mancuso lab for helpful feedback.

---

**Figure S1.**
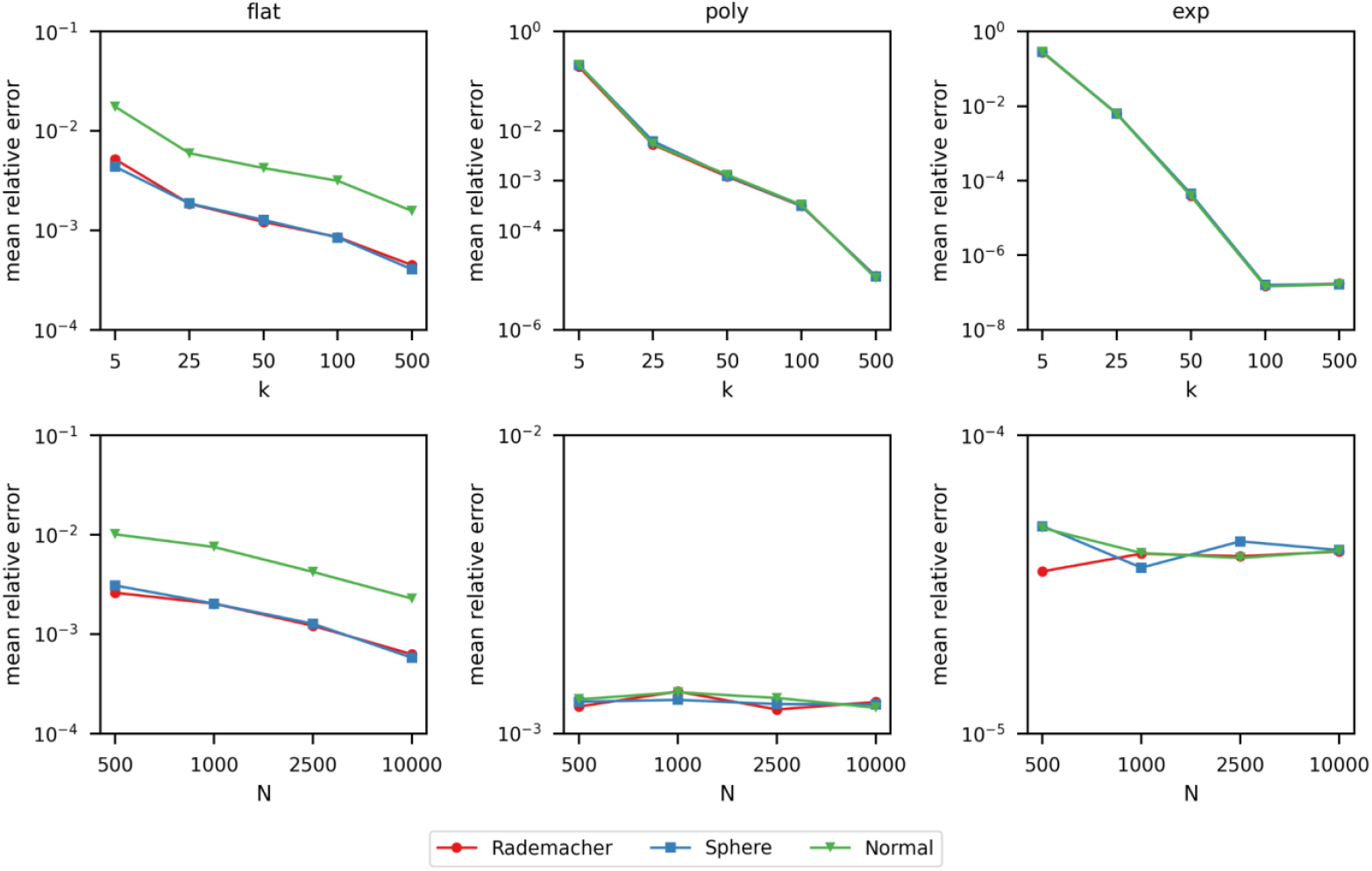
Mean relative error of XTrace with different probe distributions. *Top row:* mean relative error plotted against probe size *k*, with fixed matrix size *N*=2500. *Bottom row:* mean relative error plotted against *N* with fixed *k*=50. Columns correspond to different eigenvalue spectra: flat (left), polynomial decay (middle), and exponential decay (right). The three probe distributions compared are rademacher, sphere, and normal. Each setting was run 100 times.

**Figure S2.**
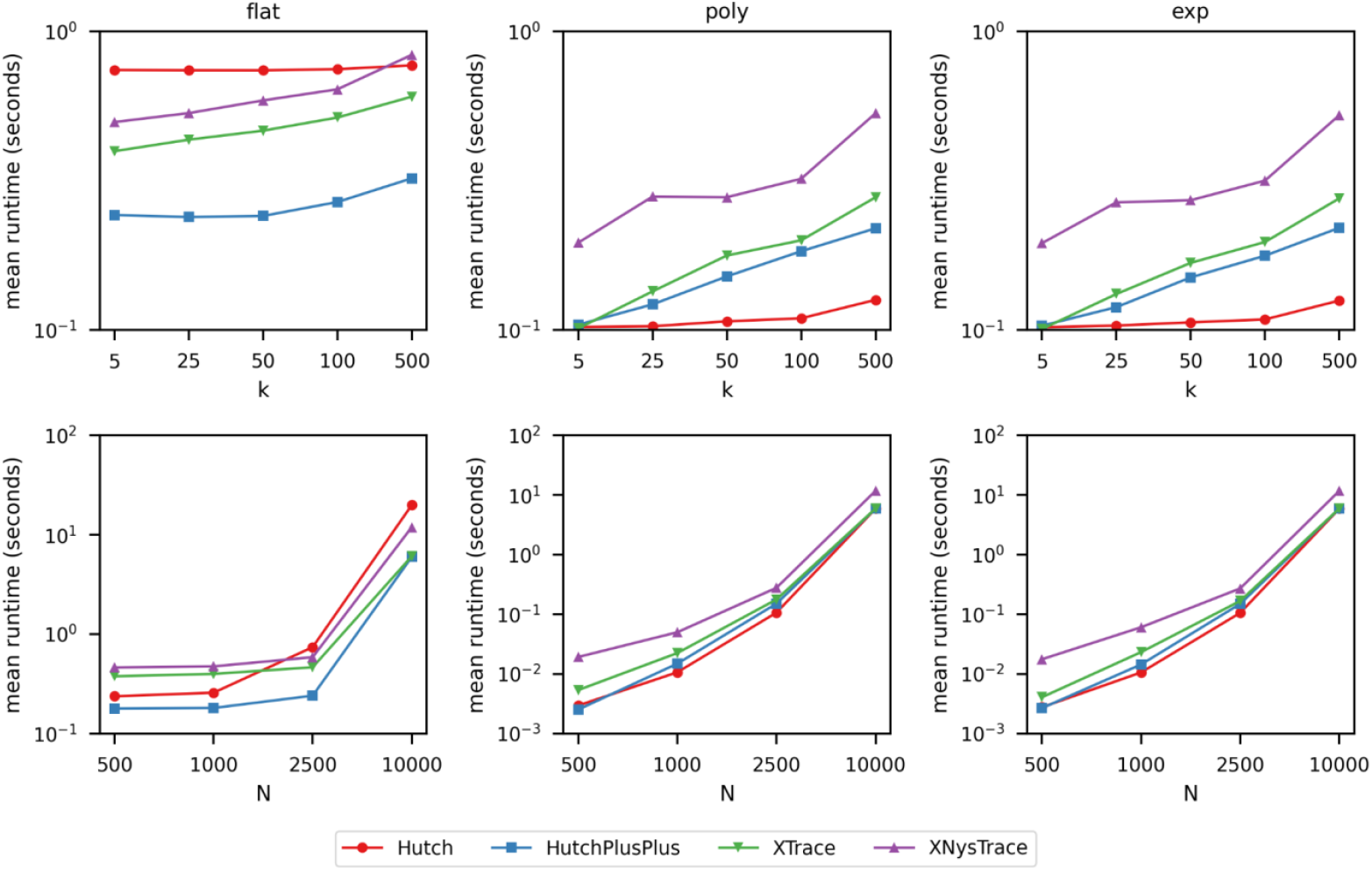
Runtime performance of different estimators. *Top row:* mean runtime plotted against probe size *k*, with fixed matrix size *N*=2500. *Bottom row:* mean runtime plotted against *N* with fixed *k*=50. Columns correspond to different eigenvalue spectra: flat (left), polynomial decay (middle), and exponential decay (right). The four methods compared are Hutch, Hutch++, XTrace, and XNysTrace. Each scenario was run 100 times.

**Figure S3.**
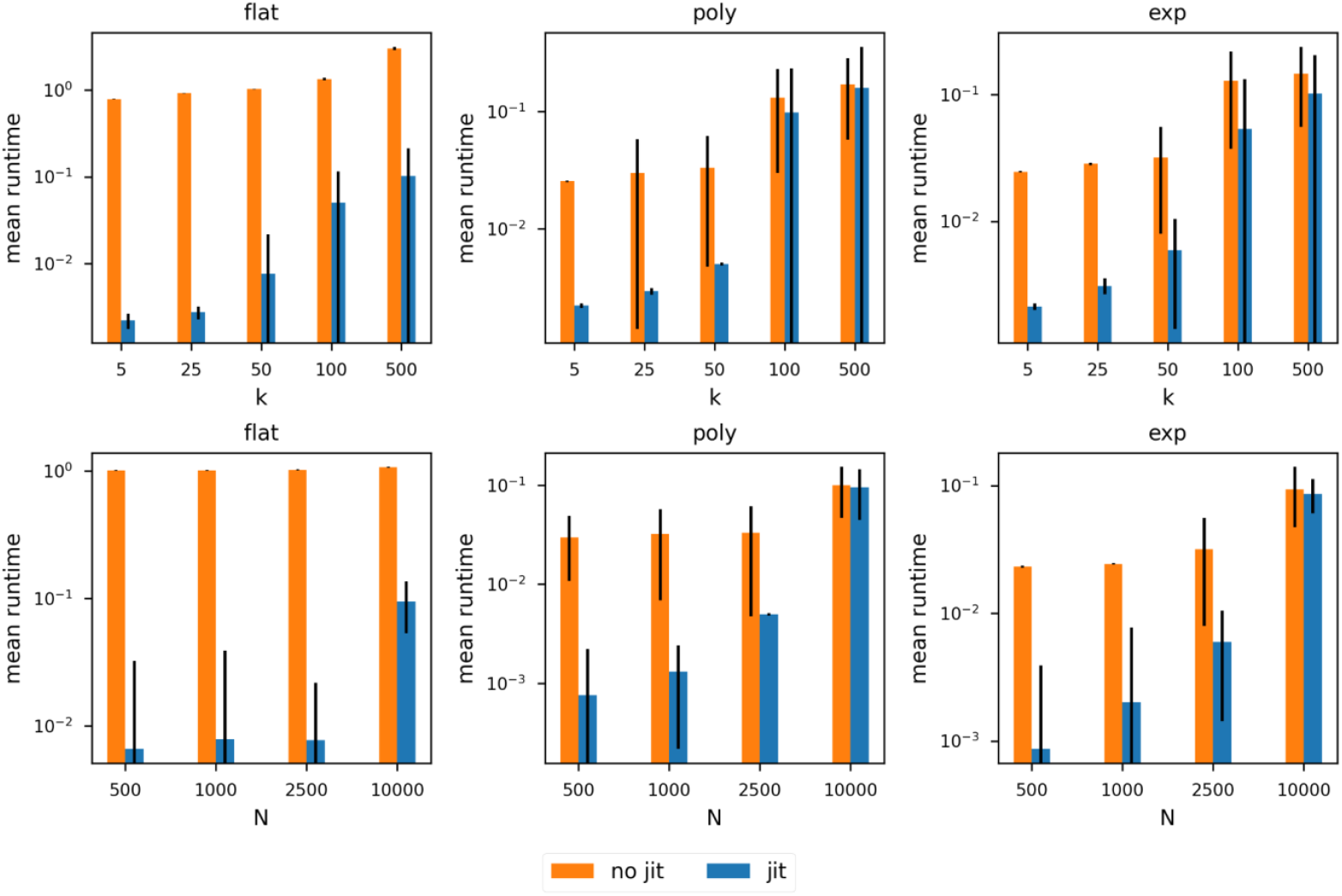
Mean runtime comparison of JIT and no-JIT version of XTrace estimation. *Top row:* mean runtime plotted against probe size *k*, with fixed matrix size *N*=2500. *Bottom row:* mean runtime plotted against *N* with fixed *k*=50. Columns correspond to different eigenvalue spectra: flat (left), polynomial decay (middle), and exponential decay (right). Orange bars indicate no JIT compilation, whereas blue bars indicate JIT compiled trace estimation. Each setting was run 100 times.

**Figure S4.**
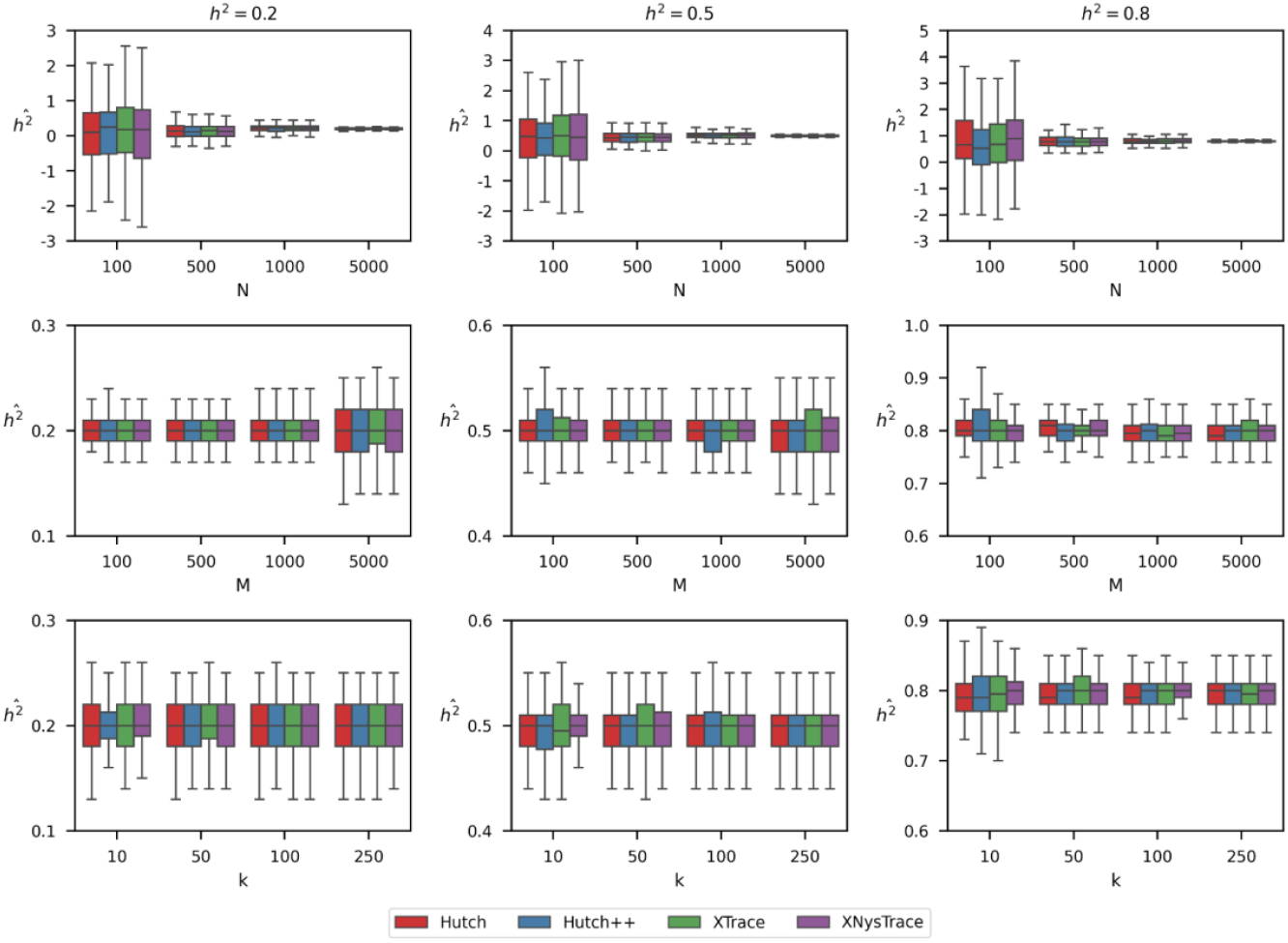
Benchmarking estimators for recovering *h*^*2*^ with the RHE-reg heritability estimator using traceax. *Top row:* ***h*^*2*^** estimates plotted against sample size *N*, with fixed number of SNPs *M*=5000 and probe size *k*=50. *Middle row:* ***h*^*2*^** estimates plotted against *M*, with fixed *N*=5000 and *k*=50. *Bottom row:* ***h*^*2*^** estimates plotted against *k*, with *N*=5000 and *M*=5000. Columns correspond to different true heritability values: ***h*^*2*^**=0.2 (left), ***h*^*2*^**=0.5 (middle), and ***h*^*2*^**=0.8 (right). The four methods compared are Hutch, Hutch++, XTrace, and XNysTrace. Each configuration was run 100 times with default parameters unless otherwise stated.

**Figure S5.**
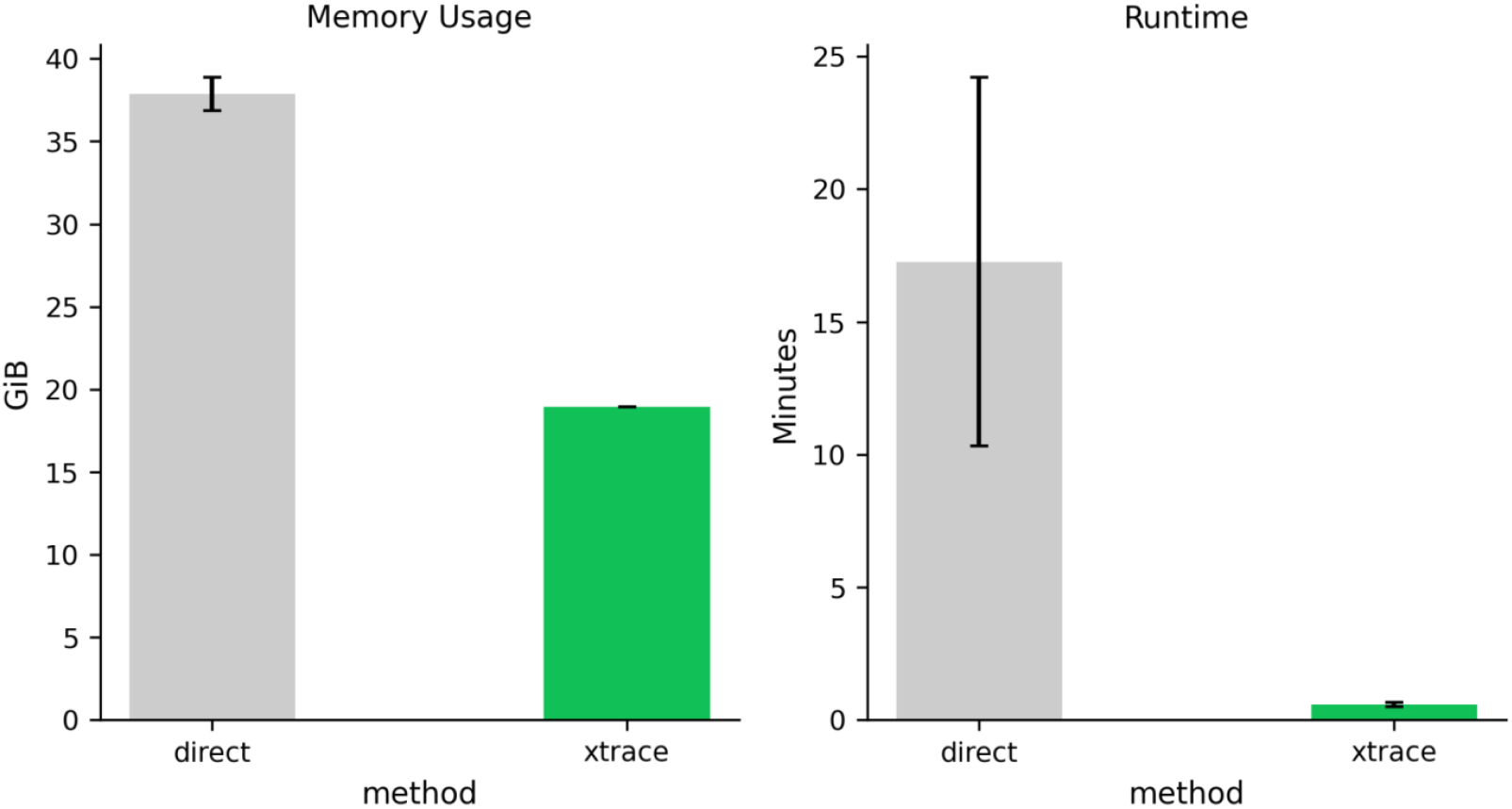
Memory usage and runtime comparison of RHE-reg SNP heritability estimation using direct vs XTrace trace computation. Across 10 independent runs, memory usage and runtime were evaluated for the RHE-reg SNP heritability estimator, comparing direct trace computation with the XTrace estimator implemented in *traceax*. Each run used *N*=50,000 individuals, *M*=50,000 SNPs, *k*=100 random probe vectors, and true heritability ***h*^*2*^**=0.5. Both methods produced accurate ***h*^*2*^** estimates.

## REFERENCES

Bradbury, J., Frostig, R., Hawkins, P., Johnson, M.J., Leary, C., Maclaurin, D., Necula, G., Paszke, A., VanderPlas, J., Wanderman-Milne, S., 2018. JAX: composable transformations of Python+ NumPy programs.

Cortinovis, A., Kressner, D., 2022. On Randomized Trace Estimates for Indefinite Matrices with an Application to Determinants. Found. Comput. Math. 22, 875–903. 10.1007/s10208-021-09525-9

Epperly, E.N., Tropp, J.A., Webber, R.J., 2024. XTrace: Making the most of every sample in stochastic trace estimation. 10.48550/arXiv.2301.07825

Gallup, O., Sechkar, K., Towers, S., Steel, H., 2024. Computational Synthetic Biology Enabled through JAX: A Showcase. ACS Synth. Biol. 13, 3046–3050. 10.1021/acssynbio.4c00307

Hutchinson, M.F., 1990. A stochastic estimator of the trace of the influence matrix for laplacian smoothing splines. Commun. Stat. - Simul. Comput. 19, 433–450. 10.1080/03610919008812866

Kaymak, M.C., Rahnamoun, A., O’Hearn, K.A., van Duin, A.C.T., Merz, K.M., Aktulga, H.M., 2022. JAX-ReaxFF: A Gradient-Based Framework for Fast Optimization of Reactive Force Fields. J. Chem. Theory Comput. 18, 5181–5194. 10.1021/acs.jctc.2c00363

Meyer, R.A., Musco, Cameron, Musco, Christopher, Woodruff, D.P., 2021. Hutch++: Optimal Stochastic Trace Estimation. 10.48550/arXiv.2010.09649

Rader, J., Lyons, T., Kidger, P., 2023. Lineax: unified linear solves and linear least-squares in JAX and Equinox. 10.48550/arXiv.2311.17283

Ubaru, S., Chen, J., Saad, Y., 2017. Fast Estimation of $tr(f(A))$ via Stochastic Lanczos Quadrature. SIAM J Matrix Anal Appl 38, 1075–1099. 10.1137/16M1104974

Ubaru, S., Saad, Y., 2018. Applications of Trace Estimation Techniques, in: Kozubek, T., Čermák, M., Tichý, P., Blaheta, R., Šístek, J., Lukáš, D., Jaroš, J. (Eds.), High Performance Computing in Science and Engineering, Lecture Notes in Computer Science. Springer International Publishing, Cham, pp. 19–33. 10.1007/978-3-319-97136-0_2

Wu, Y., Sankararaman, S., 2018. A scalable estimator of SNP heritability for biobank-scale data. Bioinforma. Oxf. Engl. 34, i187–i194. 10.1093/bioinformatics/bty253

Yeon, K., Ghosal, P., Anitescu, M., 2025. BOLT: Block-Orthonormal Lanczos for Trace estimation of matrix functions. 10.48550/arXiv.2505.12289

Zhang, Z.E., Kim, A., Suboc, N., Mancuso, N., Gazal, S., 2025. Efficient count-based models improve power and robustness for large-scale single-cell eQTL mapping. 10.1101/2025.01.18.25320755

Zvonek, J., Horning, A., Townsend, A., 2024. ContHutch++: Stochastic trace estimation for implicit integral operators. 10.48550/arXiv.2311.07035

